# Mitotic ER exit site dissociation and reassembly is regulated by TANGO1 phosphorylation status

**DOI:** 10.1101/636506

**Authors:** Miharu Maeda, Yukie Komatsu, Kota Saito

**Author notes:** Corresponding to Kota Saito.

## Abstract

Golgi fragmentation and ER exit site dissociation are considered as the leading causes of mitotic block of secretion from the ER. Although the mechanisms of Golgi fragmentation have been extensively characterized, ER exit block early in mitosis is not well-understood. We previously found that TANGO1 organizes ER exit sites by directly interacting with Sec16. Here, we showed that TANGO1 is phosphorylated by casein kinase 1 (CK1) during mitosis. Interestingly, the interaction with Sec16 was abrogated by phosphorylation of TANGO1, leading to dissociation of the ER exit sites. Moreover, a TANGO1 mutant deficient in phosphorylation inhibited the mitotic dissociation of ER exit sites. In contrast, a TANGO1 mutant mimicking CK1-mediated phosphorylation dissociated ER exit sites in interphase cells. Although CK1 activity remains constant throughout the cell cycle, PP1, a phosphatase for which activity decreases during mitosis, participates in the regulation of TANGO1 phosphorylation. This is the first report demonstrating the mechanisms of ER exit site dissociation during mitosis.

## Introduction

ER to Golgi traffic is blocked during mitosis (Kreiner and Moore, 1990; Warren et al., 1983). Mitotic Golgi are fragmented into haze, and it remains controversial whether the Golgi membrane fuses with the ER or remain segregated during mitosis (Pecot and Malhotra, 2004; Sengupta et al., 2015; Villeneuve et al., 2017; Zaal et al., 1999). The ER exit site, a specific domain on the ER for vesicle budding, dissociates during mitosis (Farmaki et al., 1999), but the mechanisms remain unclear. In interphase cells, mammalian ER exit sites consist of 2–6 cytoplasmic coat protein II (COPII)-coated pits according to electron microscopic analysis (Bannykh et al., 1996; Zeuschner et al., 2006). Although COPII-vesicle formation at ER exit sites has been extensively characterized, the biogenesis of ER exit sites is poorly understood (Maeda et al., 2019; Saito and Katada, 2015; Saito et al., 2017). Sec16, a peripheral membrane protein localized at ER exit sites, is thought to be an organizer of these sites (Miller and Schekman, 2013). However, it must first be recruited to ER membranes to perform its functions. TANGO1(L) was identified as cargo receptor for large cargoes including collagens and chylomicrons (Saito et al., 2009; Santos et al., 2016). Interestingly, we recently showed that TANGO1L, in cooperation with its alternatively-spliced isoform TANGO1S, organizes the ER exit site by interacting with and recruiting Sec16 to these sites (Maeda et al., 2017; Maeda et al., 2016). Thus, Sec16 and TANGO1 are key organizers of ER exit sites (Maeda et al., 2017; Saito and Maeda, 2019). However, it remained unknown how these organizations are regulated by the cell cycle.

ER exit sites sense alterations in signaling and change secretion profiles accordingly (Centonze and Farhan, 2019). ERK2-mediated Sec16 phosphorylation leads to an increased number of ER exit sites in response to growth factor signals (Farhan et al., 2010; Tillmann et al., 2015). In contrast, amino-acid deprivation induces Sec16 phosphorylation by ERK7, leading to ER exit site dissociation (Zacharogianni et al., 2011). Recent reports suggest that the ER contains a tyrosine kinase, LTK, which phosphorylates Sec12, a guanine-nucleotide exchange factor for Sar1, for efficient secretion (Centonze et al., 2019). Moreover, Sec24, a cargo-interacting subunit of the inner COPII-coat senses cargo folding status and activates cargo export machinery while inhibiting protein synthesis (Subramanian et al., 2019). As described, most proteins at ER exit sites have been reported to respond to external signaling; however, the mechanisms underlying ER exit site dissociation at the onset of mitosis remain poorly understood.

Here, we identified TANGO1 as a regulator of ER exit site dissociation during mitosis. TANGO1 phosphorylation was found to be regulated by kinase and phosphatase, namely CK1 and PP1, coordination. CK1-mediated TANGO1 phosphorylation reduces binding to Sec16, leading to the dissociation of ER exit sites. CK1 constantly phosphorylates TANGO1, whereas PP1 dephosphorylates TANGO1 but the activity is decreased during mitosis. Accordingly, TANGO1 phosphorylation is only upregulated during mitosis. This is the first report revealing the mechanisms of ER exit site dissociation during mitosis.

## Results

### CK1 regulates ER exit site organization

Although previous reports suggested that CK1δ is involved in COPII vesicle budding from the ER (Lord et al., 2011), the mechanisms were unclear. First, we expressed CK1α and CK1δ and evaluated their effects on ER exit site organization. CK1α expression did not affect the localization of Sec16 and Sec31 (Fig. 1A upper panel). In contrast, CK1δ expression resulted in dissociation of Sec16 and Sec31, as observed in cells depleted of TANGO1L and TANGO1S (Fig. 1A middle panel) (Maeda et al., 2017). The kinase inactive form of CK1δ, K38R, did not induce dissociation, suggesting that kinase activity is required for this process (Fig. 1A, bottom panel). Quantitative analysis showed that the colocalization efficiency between Sec16 and Sec31 was significantly reduced by expression of CK1δ but not by CK1α or CK1δ K38R (Fig. 1B). CK1δ expression did not alter the expression of other proteins tested including TANGO1 (Fig. 1C). Because the function of CK1δ considerably overlaps with CK1ε (Schittek and Sinnberg, 2014; Venerando et al., 2014), we next examined the organization of ER exit sites upon depletion of CK1δ and CK1ε. As shown in Fig. 1D, following depletion of CK1 δ and CK1ε, ER exit sites stained by Sec16, Sec23, and Sec31 were enlarged. Quantitative analysis revealed that the numbers of small and large ER exit sites were decreased and increased, respectively, following CK1 depletion (Fig. 1E), without affecting the expression of other proteins (Fig. 1F). Next, we used IC261, an ATP-competitive specific inhibitor for CK1δ and CK1ε (Mashhoon et al., 2000). Interestingly, cells treated with the inhibitor showed a similar phenotype as CK1-depleted cells (Fig. 1G and H). These results suggest that CK1δ and CK1ε are involved in organizing ER exit sites through their kinase activities. Because Sec23 has been identified as a substrate for CK1δ (Lord et al., 2011), we predicted that CK1-mediated Sec23 phosphorylation regulates the organization of ER exit sites. In this case, Sec23 depletion would eliminate the effect of IC261-mediated enlargement of ER exit sites. We depleted Sec23 from HeLa cells and examined the size of ER exit sites. As shown in Fig. S1A and B, enlarged ER exit sites were still observed following IC261 treatment, even in Sec23-depleted cells. Thus, it is unlikely that CK1-mediated regulation of ER exit site size is mediated by Sec23 phosphorylation.

**Figure 1.**
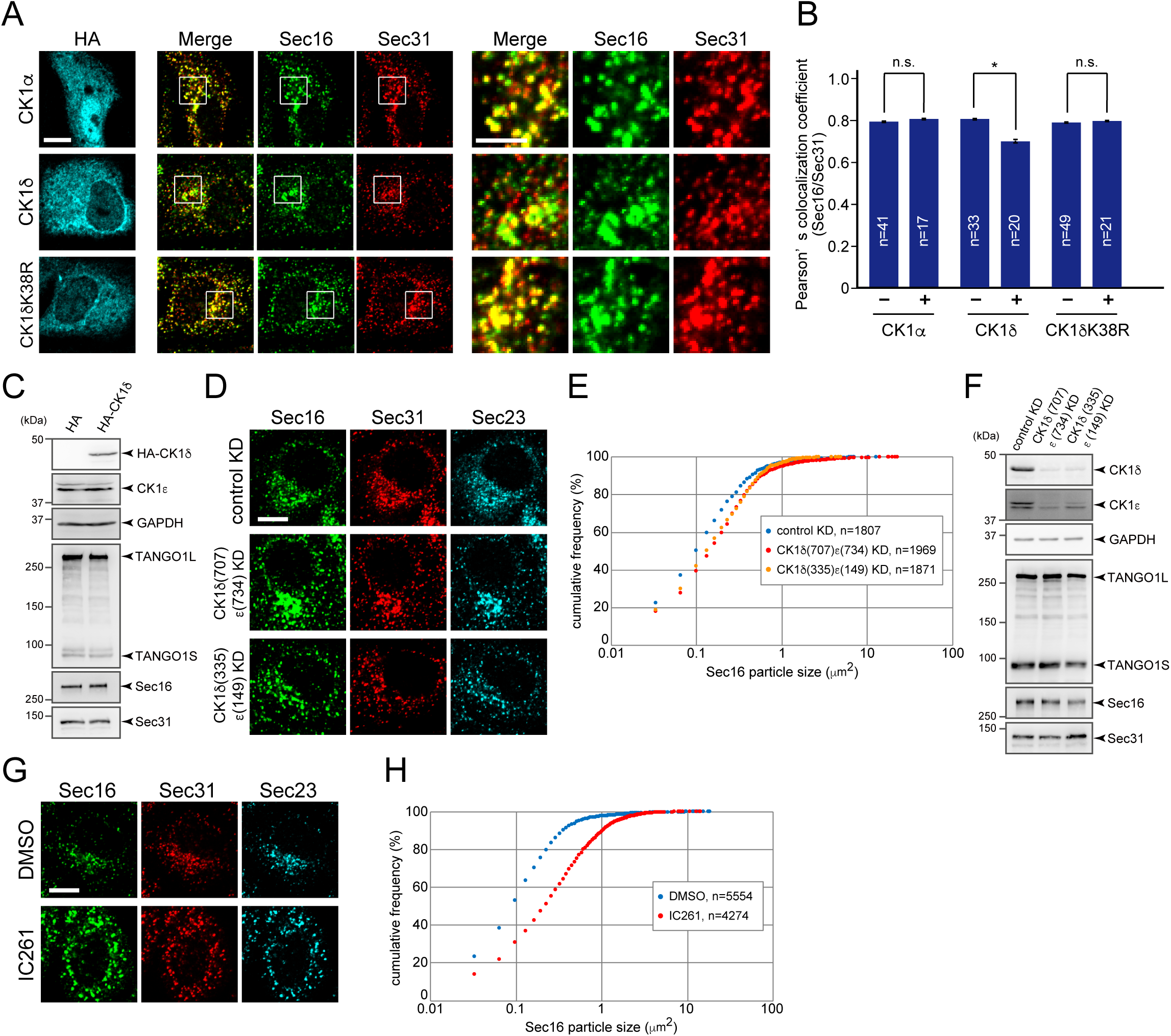
CK1 regulates ER exit site organization. (A) HeLa cells transfected with HA-CK1α, HA-CK1δ, or Ηα;−CK1δ K38R were fixed and stained with anti-Sec16-C, anti-Sec31 (m), and anti-HA antibodies. Bars = 10 *μ*m. (right) Magnification of the indicated regions on the left. Bars = 5 *μ*m. (B) Quantification of Pearson’s colocalization coefficient for (A). Error bars represent the mean ± SEM. *p < 0.05, n.s.: not significant. (C) 293T cells transfected with indicated constructs were lysed and the lysates were subjected to SDS-PAGE followed by western blotting with anti-HA, anti-CK1ε, anti-GAPDH, anti-TANGO1-CC1, anti-Sec16-N, and anti-Sec31 (r) antibodies. (D) HeLa cells transfected with the indicated siRNAs were fixed and stained with anti-Sec16-C, anti-Sec31(m) and anti-Sec23 antibodies. Bars = 10 *μ*m. (E) Quantification of ER exit site size distribution from (D). (F) HeLa cells transfected with indicated siRNAs were lysed and the lysates were subjected to SDS-PAGE followed by western blotting with anti-CK1δ, anti-CK1ε, anti-GAPDH, anti-TANGO1-CC1, anti-Sec16-N, and anti-Sec31(r) antibodies. (G) HeLa cells were treated with either DMSO or 10 *μ*M of IC261 for 12 h. Cells were fixed and stained with anti-Sec16-C, anti-Sec31(m), and anti-Sec23 antibodies. Bars = 10 *μ*m. (H) Quantification of ER exit site size distribution from (G).

### CK1 reduces the interaction between Sec16 and TANGO1

To further investigate the effect of CK1 on ER exit site organization, we determined whether CK1 interacts with other proteins at ER exit sites. As shown in Fig. 2A, we found that CK1δ interacts with Sec16 (Fig. 2A, lanes 1 and 2). The kinase activity of CK1δ did not affect this interaction (Fig. 2A, lanes 3 and 4). To determine the interaction domain on Sec16, we examined interactions between Sec16 deletion constructs and CK1 (Fig. 2B). Sec16 (1101–1600) and Sec16 (1601–1890) efficiently interacted with CK1δ (Fig. 2C). When Sec16 was truncated into the TANGO1-interacting region (1101–1400), the interaction was reduced but not eliminated (Fig. 2C) (Maeda et al., 2017). These results suggest that Sec16 possesses multiple CK1-interacting regions that can be mapped to 1101–1890. In contrast, TANGO1S failed to interact with CK1 (Fig. 2D). We previously showed that the interaction between TANGO1 and Sec16 is vital for proper ER exit site organization (Maeda et al., 2017). Thus, we next examined whether CK1 regulates ER exit site organization via TANGO1/Sec16. Our previous data showed that TANGO1S and TANGO1L have redundant functions in organizing ER exit sites; thus, we used only TANGO1S for further assays (Maeda et al., 2017). As expected, Sec16 interacted with both TANGO1S and CK1δ; however, the interaction between TANGO1 and Sec16 was markedly reduced when CK1δ was added (Fig. 2E). These results indicate that CK1 organizes ER exit sites by modulating TANGO1–Sec16 interactions.

**Figure 2.**
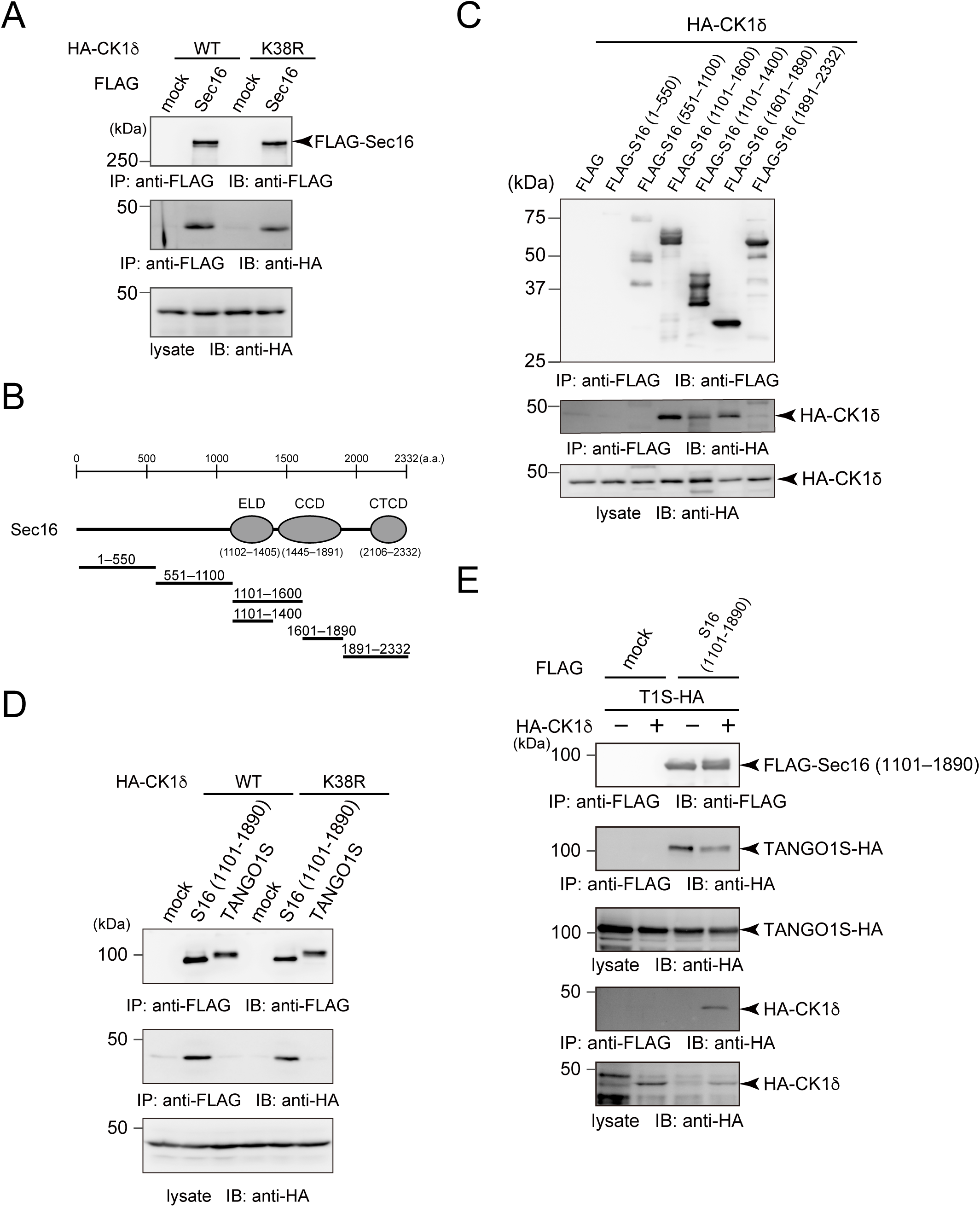
CK1 reduces the interaction between Sec16 and TANGO1. (A) 293T cells were transfected with HA-CK1δ or HA-CK1δ K38R with FLAG-Sec16 constructs as indicated. Cell lysates were immunoprecipitated with anti-FLAG antibody and eluted with a FLAG peptide. Eluates were then subjected to SDS-PAGE followed by western blotting with anti-FLAG and anti-HA antibodies. (B) Schematic representation of human Sec16 domain organization. ELD, ER exit site localization domain; CCD, central conserved domain; CTCD, C-terminal conserved domain. (C) 293T cells were transfected with indicated FLAG-tagged Sec16 deletion constructs with HA-CK1δ constructs as indicated. Cell lysates were immunoprecipitated with anti-FLAG antibody and eluted with a FLAG peptide. Eluates were then subjected to SDS-PAGE followed by western blotting with anti-FLAG and anti-HA antibodies. (D) 293T cells were transfected with FLAG-Sec16 (1101–1890 aa) or TANGO1S-FLAG and HA-CK1δ or HA-CK1δ K38R constructs. Cell lysates were immunoprecipitated with anti-FLAG antibody and eluted with a FLAG peptide. Eluates were then subjected to SDS-PAGE followed by western blotting with anti-FLAG and anti-HA antibodies. (E) 293T cells were transfected with FLAG-Sec16 (1101–1890 aa), TANGO1S-HA, and HA-CK1δ constructs as indicated. Cell lysates were immunoprecipitated with anti-FLAG antibody and eluted with a FLAG peptide. Eluates were then subjected to SDS-PAGE followed by western blotting with anti-FLAG and anti-HA antibodies.

### CK1-mediated TANGO1 phosphorylation dissociates ER exit sites

The above results suggest that the kinase activity of CK1 is important for ER exit site organization. Thus, we searched for possible phosphorylation sites within TANGO1 and Sec16 using the PhosphositePlus and NetPhos databases (Blom et al., 1999; Hornbeck et al., 2015). We found phosphorylation hotspots within TANGO1 just upstream of the proline-rich domain (PRD) and named these sites as phosphorylation predicted sequences (PPS) (Fig. 3A). PPS alignment indicated that this region is well-conserved among species (Fig. 3B). Next, we mutated serine or threonine to alanine in the PPS (SA mutant) and evaluated whether this region is phosphorylated via CK1 in an *in vitro* kinase assay. We used CK1 without the C-terminal regulatory domain for hyperactivation (Graves and Roach, 1995). Recombinant TANGO1 PPS+PRD or the corresponding SA mutant were incubated either with wild-type or kinase-dead CK1δ in the presence of [γ-^32^P] ATP (Fig. S2). As shown in Fig. 3C, wild-type CK1δ efficiently phosphorylated TANGO1 PPS+PRD but failed to phosphorylate the SA mutant, strongly suggesting that the TANGO1 PPS is phosphorylated by CK1δ. The inefficient phosphorylation by kinase-dead CK1 validated the specificity of this effect. Next, we prepared TANGO1S with the phosphorylation mimic (TANGO1S (SE)-HA) or inefficient mutations (TANGO1S (SA)-HA) and examined their interactions with Sec16. Coimmunoprecipitation analysis revealed that SE mutants reduced the binding to Sec16, whereas SA mutants enhanced the binding to Sec16 compared to wild-type TANGO1S (Fig. 4A). These data indicate that phosphorylation of TANGO1 by CK1 reduced the interaction with Sec16. Indeed, the interaction between TANGO1 and Sec16 was not susceptible to CK1δ when the TANGO1 SA mutant was used rather than the wild-type protein (Compare Fig. 2E and Fig. 4B). Next, we examined whether TANGO1 mutants affect the organization of ER exit sites. We previously showed that Sec16 and Sec31 dissociates upon TANGO1 depletion, which cannot be rescued by TANGO1 lacking the Sec16-interacting region (Maeda et al., 2017). We depleted TANGO1L and TANGO1S and expressed TANGO1 SE and SA mutants. Immunofluorescence data are shown in Fig. 4C and quantitative data in Fig. 4D. Although wild-type TANGO1S and SA mutants efficiently rescued the dissociation between Sec16 and Sec31, SE mutants failed to correct the dissociation of ER exit sites. These data strongly suggest that CK1-mediated TANGO1 phosphorylation in PPS regulates ER exit site organization. We made series of mutations in the TANGO1 PPS to identify the critical residues for CK1 phosphorylation (Fig. S3A). We examined not only the binding properties between Sec16 and TANGO1 mutants (Fig. S3B), but also the organizing efficiency of TANGO1 mutants (Fig. S3C). The data indicated that the number of mutations affects both the binding properties and organization, while critical residues important for regulating ER exit site organization were not likely existed. Thus, the degree of phosphorylation in TANGO1’s PPS by CK1 regulates the organization of ER exit sites.

**Figure 3.**
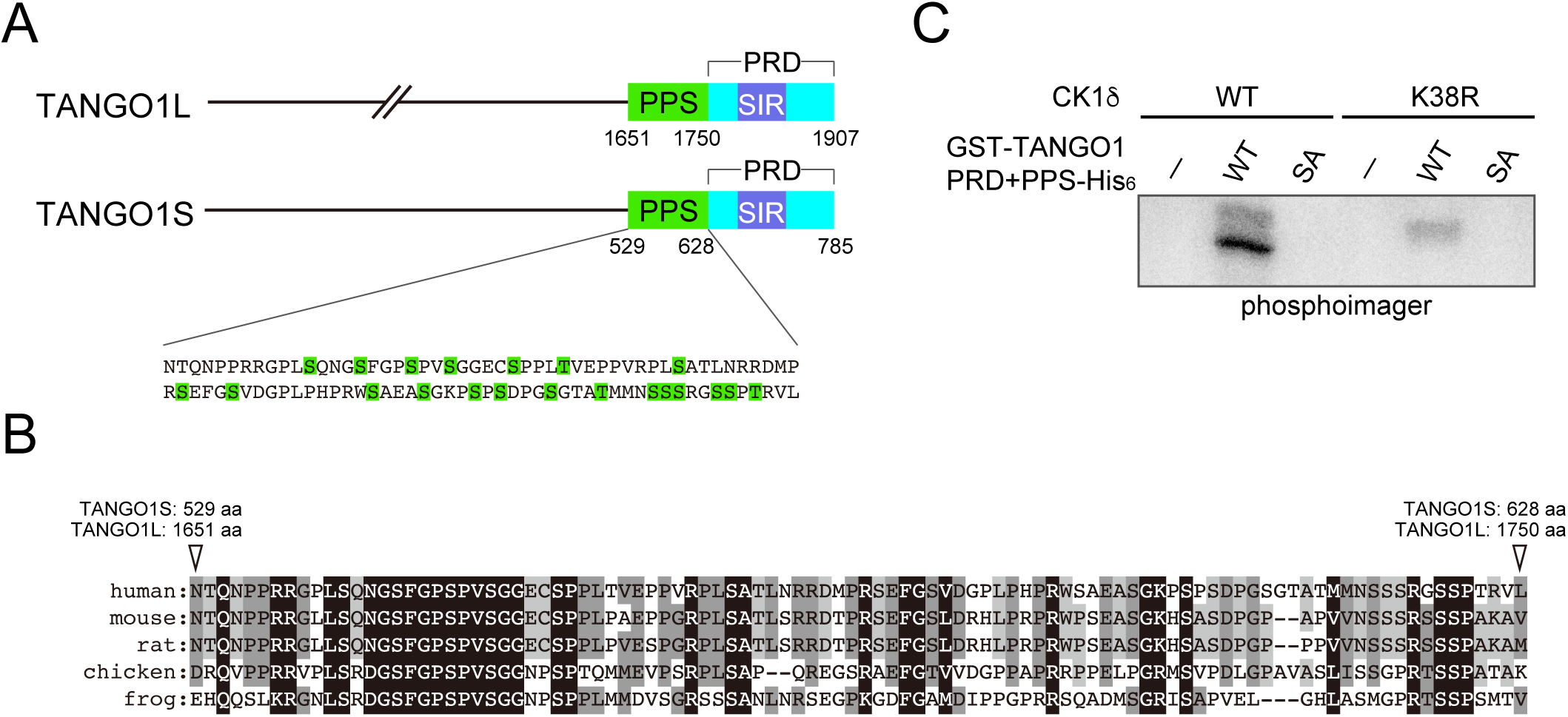
The PPS domain of TANGO1 is phosphorylated by CK1 in an *in vitro* kinase assay. (A) Schematic representation of TANGO1 domain organization. PPS, phosphorylation predicted sequences; SIR, Sec16-interacting region; PRD, proline-rich domain. (B) Alignment of human TANGO1 PPS with corresponding regions from other species. Identical amino acids are shaded in black and similar amino acids are shaded in gray. (C) *In vitro* kinase assay was performed with purified recombinant GST-TANGO1 PPS+PRD-His_6_ or GST-TANGO1 PPS+PRD (SA)-His_6_ with CK1δ (1–317 aa).

**Figure 4.**
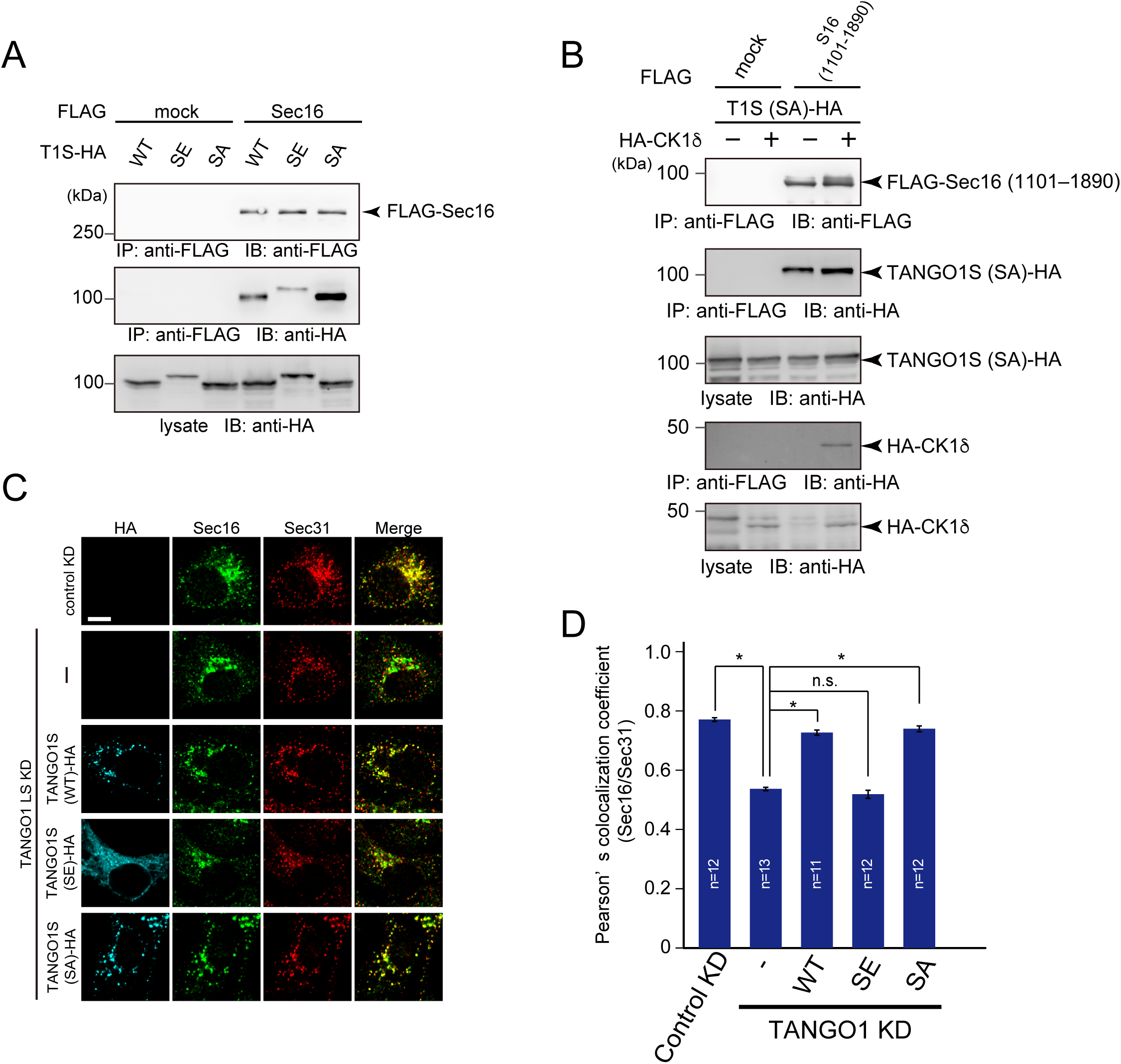
CK1-mediated TANGO1 phosphorylation dissociates ER exit sites. (A) 293T cells were transfected with FLAG-Sec16 and TANGO1S-HA or TANGO1S (SE)- HA or TANGO1S (SA)-HA constructs. Cell lysates were immunoprecipitated with an anti-FLAG antibody and eluted with a FLAG peptide. Eluates were then subjected to SDS-PAGE followed by western blotting with anti-FLAG and anti-HA antibodies. (B) 293T cells were transfected with FLAG-Sec16 (1101–1890 aa), TANGO1S (SA)-HA, and HA-CK1δ constructs as indicated. Cell lysates were immunoprecipitated with an anti-FLAG antibody and eluted with a FLAG peptide. Eluates were then subjected to SDS-PAGE followed by western blotting with anti-FLAG and anti-HA antibodies. (C) HeLa cells were treated with TANGO1L and TANGO1S siRNAs. After 24 h, indicated HA-tagged TANGO1S constructs were transfected and cells were further cultured for 24 h. The cells were fixed and stained with anti-Sec16, anti-Sec31, and anti-HA antibodies. Bars = 10 *μ*m. (D) Quantification of Pearson’s colocalization coefficient for (C). Error bars represent means ± SEM. *p < 0.05, n.s.: not significant.

### Mitotic ER exit site dissociation is mediated by TANGO1 phosphorylation

Secretion from the ER is blocked during mitosis, which dramatically affects ER exit site organization (Farmaki et al., 1999; Kreiner and Moore, 1990). However, the mechanisms of ER exit site dissociation during mitosis are unknown. Previous reports suggest that Sec16 and Sec31 are disorganized during mitosis (Hughes and Stephens, 2010). These results prompted us to investigate whether TANGO1 phosphorylation is involved in ER exit site disorganization during mitosis. Cells were synchronized with double thymidine block, and metaphase cells were selected to analyze ER exit sites. As shown in Fig. 5A, Sec16 and Sec31 showed distinct staining in metaphase, as previously reported (Hughes and Stephens, 2010). Quantitative analysis clearly indicated that colocalization efficiency between Sec16 and Sec31 was significantly reduced during mitosis (Fig. 5B). Notably, reduced colocalization was also observed between TANGO1 and Sec31 in mitotic cells, suggesting that ER exit site dissociation occurs for both peripheral membrane and membrane proteins (Fig. 5C and D).

**Figure 5.**
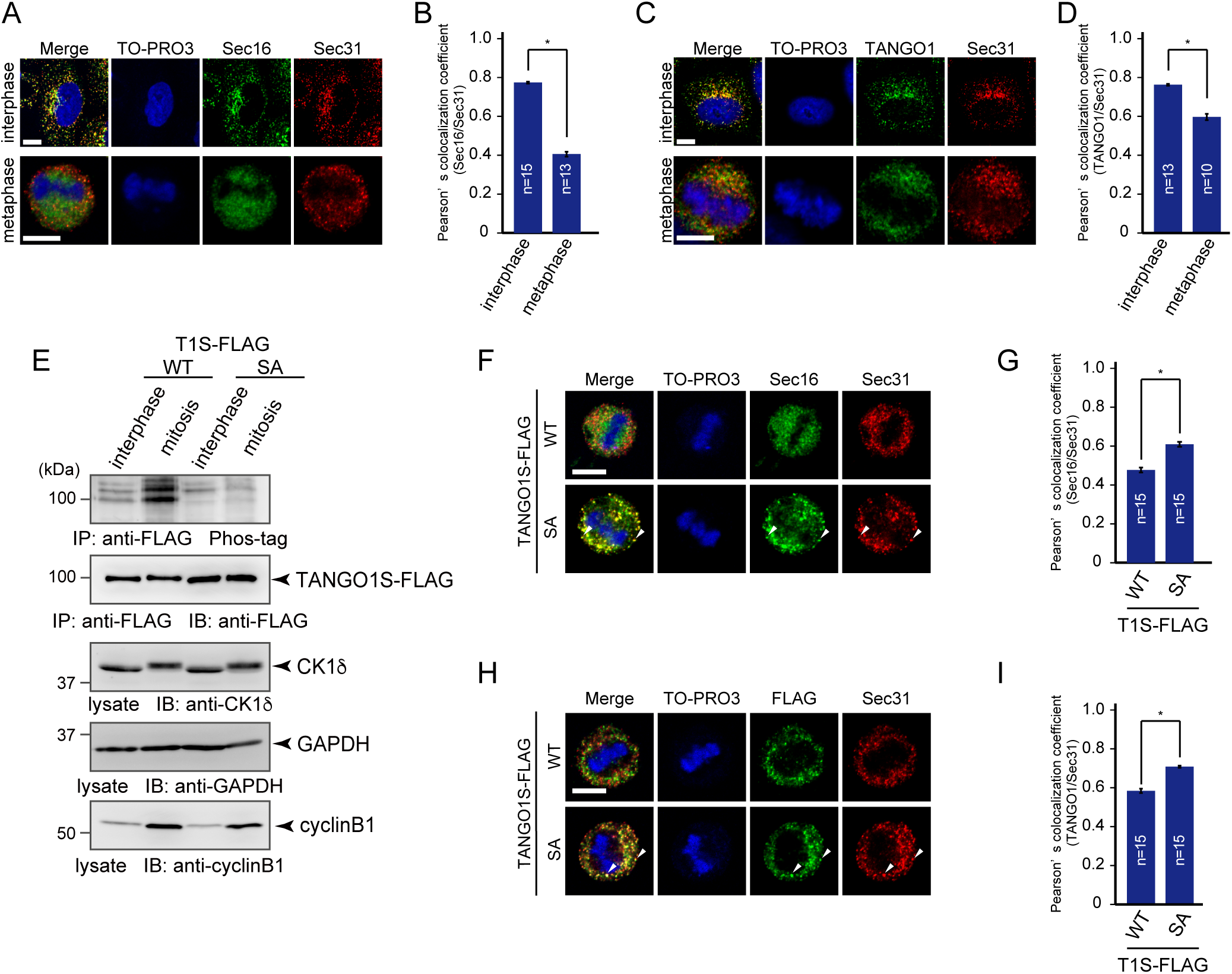
The phosphorylation of TANGO1 is required for mitotic ER exit site dissociation. (A) Non-treated HeLa cells and HeLa cells synchronized at mitotic phase with double thymidine block were fixed and stained with anti-Sec16-C, anti-Sec31(m), or TO-PRO3. Bars = 10 *μ*m. (B) Quantification of Pearson’s colocalization coefficient for (A). (C) Non-treated HeLa cells and HeLa cells synchronized at mitotic phase with double thymidine block were fixed and stained with anti-TANGO1-CT, anti-Sec31(m), or TO-PRO3. Bars = 10 *μ*m. (D) Quantification of Pearson’s colocalization coefficient for (C). (E) Interphase extracts of doxycycline-inducible HeLa cells expressing either TANGO1S-FLAG or TANGO1S (SA)-FLAG were collected. Mitotic extracts were collected through treatment of nocodazole followed by shake-off. Cell lysates were immunoprecipitated with an anti-FLAG antibody and eluted with a FLAG peptide. Eluates were subjected to SDS-PAGE followed by biotinylated Phos-tag detection or western blotting with an anti-FLAG antibody. Lysates were subjected to SDS-PAGE followed by western blotting with anti-CK1δ, anti-GAPDH, and anti-cyclinB1 antibodies. (F) Doxycycline-inducible HeLa cells expressing either TANGO1S-FLAG or TANGO1S (SA)-FLAG were synchronized in the mitotic phase by double thymidine block. Cells were then fixed and stained with anti-Sec16-C, anti-Sec31(m), or TO-PRO3. Bars = 10 *μ*m. (G) Quantification of Pearson’s colocalization coefficient for (F). (H) Doxycycline-inducible HeLa cells expressing either TANGO1S-FLAG or TANGO1S (SA)-FLAG were synchronized in the mitotic phase by double thymidine block. Cells were then fixed and stained with anti-FLAG, anti-Sec31 (m), and TO-PRO3. Bars = 10 *μ*m. (I) Quantification of Pearson’s colocalization coefficient for (H). Error bars represent the means ± SEM. *p < 0.05, n.s.: not significant.

We thus evaluated whether TANGO1 phosphorylation is upregulated during mitosis. Phos-tag blot analysis of cells stably expressing TANGO1 constructs revealed that wild-type TANGO1 is hyperphosphorylated during mitosis (Fig. 5E). In contrast, TANGO1 SA mutants were not phosphorylated in mitotic cells, suggesting that mitotic TANGO1 phosphorylation predominantly occurs at the PPS (Fig. 5E). Based on these results, we hypothesized that ER exit site dissociation during mitosis is regulated by TANGO1 phosphorylation.

To test this, we investigated if the expression of TANGO1 SA mutants affects ER exit site organization during mitosis. Cells stably expressing wild-type TANGO1 or TANGO1 SA mutants were synchronized at metaphase and Sec16/Sec31 and TANGO1/Sec31 colocalization rates were quantified. As shown in Fig. 5F and H, metaphase cells expressing wild-type TANGO1 showed reduced Sec16/Sec31 and TANGO1/Sec31 colocalization rates compared to those in wild-type cells. Conversely, metaphase cells expressing TANGO1 SA mutants clearly showed punctate structures that were stained both by Sec16/Sec31 and TANGO1/Sec31. Quantitative analysis suggested that cells expressing TANGO1 SA mutants had significantly increased colocalization efficiencies between two components (Fig. 5 G and I). These results strongly suggest that mitotic ER exit site dissociation is mediated by TANGO1 hyperphosphorylation during mitosis.

### PP1 is involved in the TANGO1-mediated ER exit site organization

Evidence that CK1 activity is enhanced during mitosis is lacking. Thus, we speculated that other kinases or phosphatases are involved in TANGO1 hyperphosphorylation during mitosis. It has been known that protein phosphatase1 (PP1) and protein phosphatase 2A (PP2A)-B55 activities are downregulated during mitosis (Nilsson, 2019). Thus, we checked if these phosphatases are involved in ER exit site organization. Although endothall, a specific inhibitor of PP2, had no effects on ER exit site organization, cells treated with okadaic acid (PP1/PP2 inhibitor) showed reduced colocalization efficiency between Sec16 and Sec31 (Fig. 6A and B). The depletion of PP1 by siRNAs also reduced colocalization between Sec16 and Sec31, strongly suggesting that PP1 is involved in ER exit site organization (Fig. 6C–E). We then checked the phosphorylation status of TANGO1 upon okadaic-acid treatment. TANGO1 phosphorylation was significantly increased by okadaic acid (Fig. 6F), indicating that PP1 dephosphorylates TANGO1 in interphase cells. In addition, TANGO1 SA mutants showed no changes upon drug treatment (Fig. 6F). These data suggest that PP1 predominantly dephosphorylates the PPS region of TANGO1, which can be phosphorylated by CK1. We next checked whether ER exit site dissociation mediated by okadaic acid treatment could be rescued by the expression of TANGO1 SA mutants. As shown in Fig. 6G, the expression of TANGO1 SA mutants significantly increased colocalization efficiency between Sec16 and Sec31 under okadaic acid treatment. Together, ER exit site organization is regulated by TANGO1 phosphorylation status, which can be controlled by CK1-mediated phosphorylation and PP1-mediated dephosphorylation.

**Figure 6.**
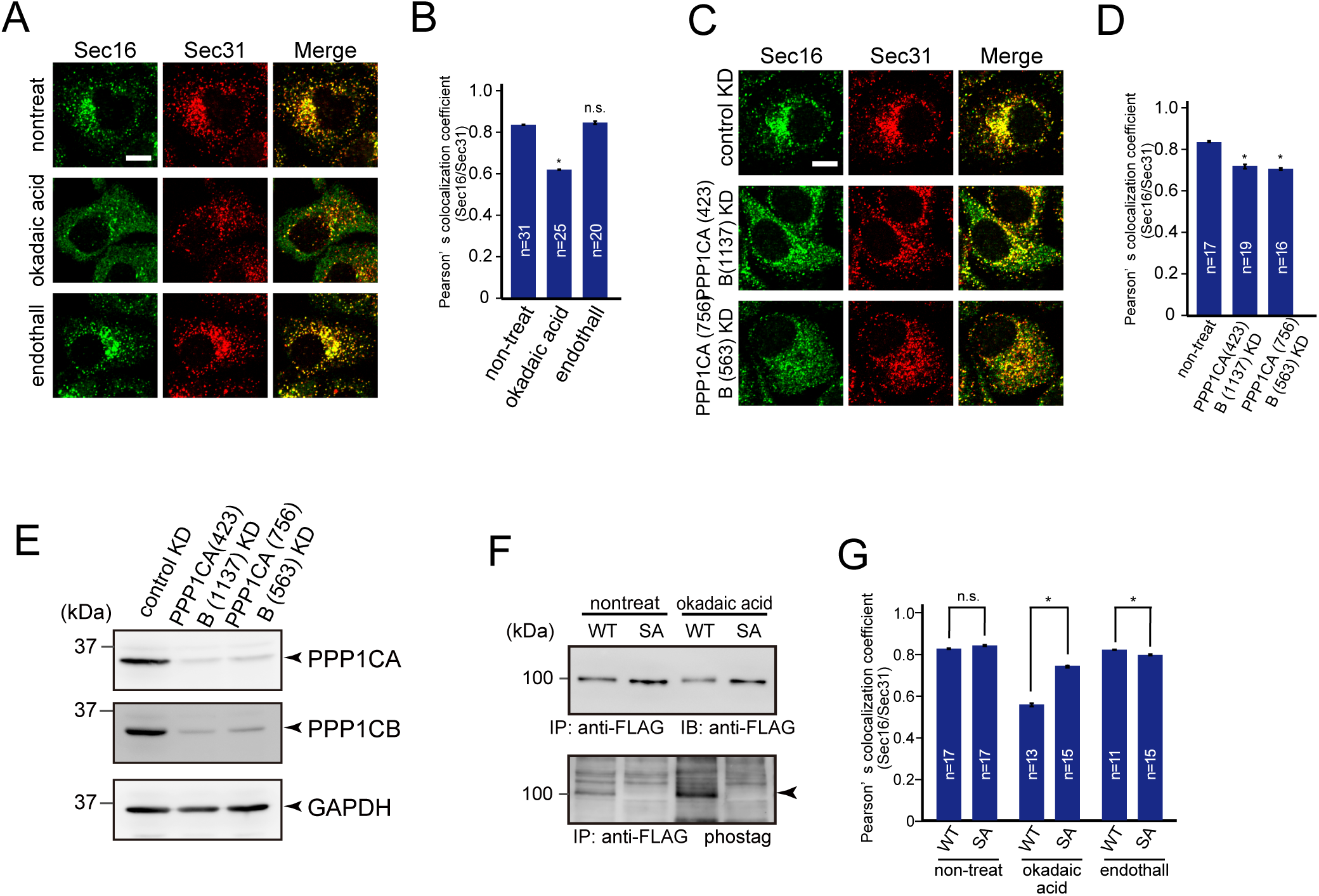
Inhibition of PP1 results in ER exit site dissociation. (A) HeLa cells were not treated or treated with 50 nM okadaic acid or 1 *μ*M endothall for 2 h. Cells were fixed and stained with anti-Sec16-C and anti-Sec31(m) antibodies. Bars = 10 *μ*m. (B) Quantification of Pearson’s colocalization coefficient for (A). (C) HeLa cells transfected with the indicated siRNAs were fixed and stained with anti-Sec16-C and anti-Sec31(m) antibodies. Bars = 10 *μ*m. (D) Quantification of Pearson’s colocalization coefficient for (C). (E) HeLa cells transfected with indicated siRNAs were lysed and the lysates were subjected to SDS-PAGE followed by western blotting with anti-PPP1CA, anti-PPP1CB, and anti-GAPDH antibodies. (F) Doxycycline-inducible HeLa cells expressing either TANGO1S-FLAG or TANGO1S (SA)-FLAG were not treated or treated with 50 nM okadaic acid for 2 h. Cell lysates were immunoprecipitated with an anti-FLAG antibody and eluted with a FLAG peptide. Eluates were subjected to SDS-PAGE followed by biotinylated Phos-tag detection or western blotting with an anti-FLAG antibody. (G) Doxycycline-inducible HeLa cells expressing either TANGO1S-FLAG or TANGO1S (SA)-FLAG were not treated or treated with 50 nM okadaic acid or 1 *μ*M endothall for 2 h. Cells were fixed and stained with anti-Sec16-C and anti-Sec31(m) antibodies and quantified based on Pearson’s colocalization coefficient. Error bars represent the mean ± SEM. *p < 0.05, n.s.: not significant.

### CK1 is required for mitotic ER exit site dissociation

PP1 activity is repressed through phosphorylation by Cdk1–cyclinB1 from the early onset of mitosis to anaphase. If CK1 is the predominant kinase for TANGO1 phosphorylation and TANGO1 hyperphosphorylation is due to the reduced kinase activity of PP1 during mitosis, CK1 depletion would also inhibit ER exit site dissociation during mitosis. As shown in Fig. 7A, CK1 δ/ε-depleted cells synchronized at metaphase clearly showed punctate structures that were stained by both Sec16 and Sec31. Quantitative analysis suggested that cells depleted of CK1 δ/ε showed significantly increased colocalization efficiency between Sec16 and Sec31 (Fig. 7B). Cells synchronized at prometaphase by nocodazole block also showed punctate structures visualized by detecting Sec16 and Sec31 upon CK1 δ/ε knockdown (Fig. S4A and B). Moreover, colocalization between TANGO1 and Sec31 was also increased during mitosis upon CK1 δ/ε depletion (Fig. 7C and D). These data suggest that CK1 δ/ε are the predominant kinases mediating TANGO1 phosphorylation for mitotic ER exit site dissociation.

**Figure 7.**
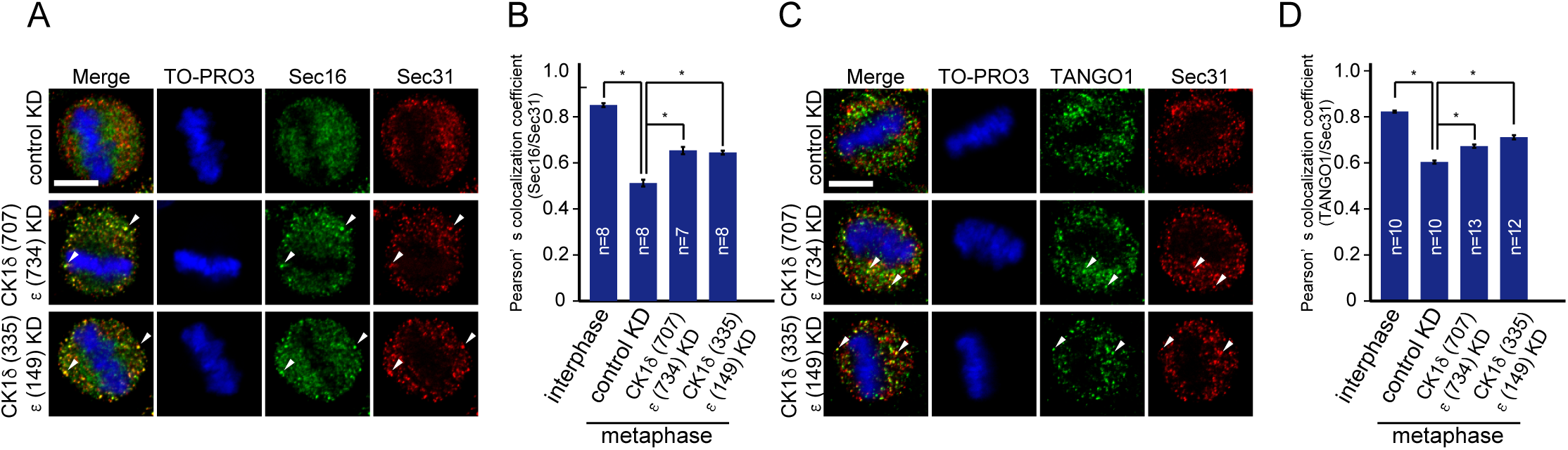
CK1 is required for mitotic ER exit site dissociation. (A) HeLa cells transfected with indicated siRNAs were synchronized at mitotic phase with double thymidine block. Cells were then fixed and stained with anti-Sec16-C, anti-Sec31(m), or TO-PRO3. Bars = 10 *μ*m. (B) Quantification of Pearson’s colocalization coefficient for (A). (C) HeLa cells transfected with indicated siRNAs were synchronized at mitotic phase with double thymidine block. Cells were then fixed and stained with anti-TANGO1-CT, anti-Sec31(m), or TO-PRO3. Bars = 10 *μ*m. (D) Quantification of Pearson’s colocalization coefficient for (C). Error bars represent the mean ± SEM. *p < 0.05.

## Discussion

We previously showed that TANGO1 and Sec16 organize ER exit sites through their interaction (Maeda et al., 2017). In this study, we identified TANGO1 as a regulator of ER exit site organization during the cell cycle. We showed that the TANGO1 PPS is hyperphosphorylated during mitosis, which reduces binding affinity to Sec16, leading to the dissociation of ER exit sites. Interestingly, the interaction domain of TANGO1 for Sec16 is not on the PPS but rather on the Sec16-interacting region, a domain approximately 40 amino acids downstream of the PPS. Structural analysis is necessary to determine how PPS phosphorylation affects the binding between TANGO1 and Sec16. We tried to map the critical phosphorylation residues of TANGO1 PPS; however, it seems likely that multiple phosphorylation events occur in the PPS and the degree of phosphorylation correlates with the binding affinity for Sec16. Similar regulation was observed for the NDC80 complex, in which multiple phosphorylation events change the affinity for microtubules (Zaytsev et al., 2015). These results imply that a balance of kinase and phosphatase activities is crucial to regulate multiple TANGO1 phosphorylation events, which then regulates the organization of ER exit sites (Gelens et al., 2018).

During mitosis, intracellular organelles and domains undergo extensive rearrangements to achieve controlled division (Jongsma et al., 2015) (Fig. 8A). In addition, increased evidence suggest that intracellular membrane trafficking is attenuated during mitosis (Yeong, 2013). Mechanisms of Golgi inheritance have been extensively characterized, and a considerable number of kinases including Cdk1, Mek1, Erk1/2, Myt1, and Plk1 were identified as responsible for Golgi fragmentation at the entry of mitosis (Valente and Colanzi, 2015). Kinase activity has also been shown to be important for diminished traffic within the Golgi (Nakamura et al., 1997; Stuart et al., 1993). It is still controversial though whether fragmented Golgi stays as a Golgi haze or fuses with the ER during metaphase (Sengupta et al., 2015; Villeneuve et al., 2017) (Fig. 8A). In either case, Golgi reassembly begins at telophase. Agreement has yet to be reached as to how the ER divides during mitosis. The ER cisternae were reported to be remodeled into tubules, although other reports suggested that they remain throughout the cell cycle (Lu et al., 2009; Puhka et al., 2007). In addition, it has been shown that export from the ER is attenuated (Kreiner and Moore, 1990; Warren et al., 1983) and ER exit site are dissociated during mitosis (Hammond and Glick, 2000; Hughes and Stephens, 2010). However, compared to the number of kinases identified during Golgi fragmentation, few have been reported to regulate ER exit site organization and ER export during mitosis. Cdc2-mediated p47 phosphorylation was reported to disorganize ER exit sites in semi-intact cells, but how this signal regulates the ER exit site remains unknown (Kano et al., 2004). Moreover, a further study suggested that p47 phosphorylation rather regulates ER network disruption, indicating that the effect of p47 on the ER exit site might be indirect (Kano et al., 2005). In this study, we identified CK1 and PP1 as a kinase and a phosphatase that directly regulate ER exit site organization via TANGO1 during mitosis. We propose that CK1-mediated disassembly of ER exit sites would be a critical event regulating the mitotic block in secretion from the ER.

**Figure 8.**
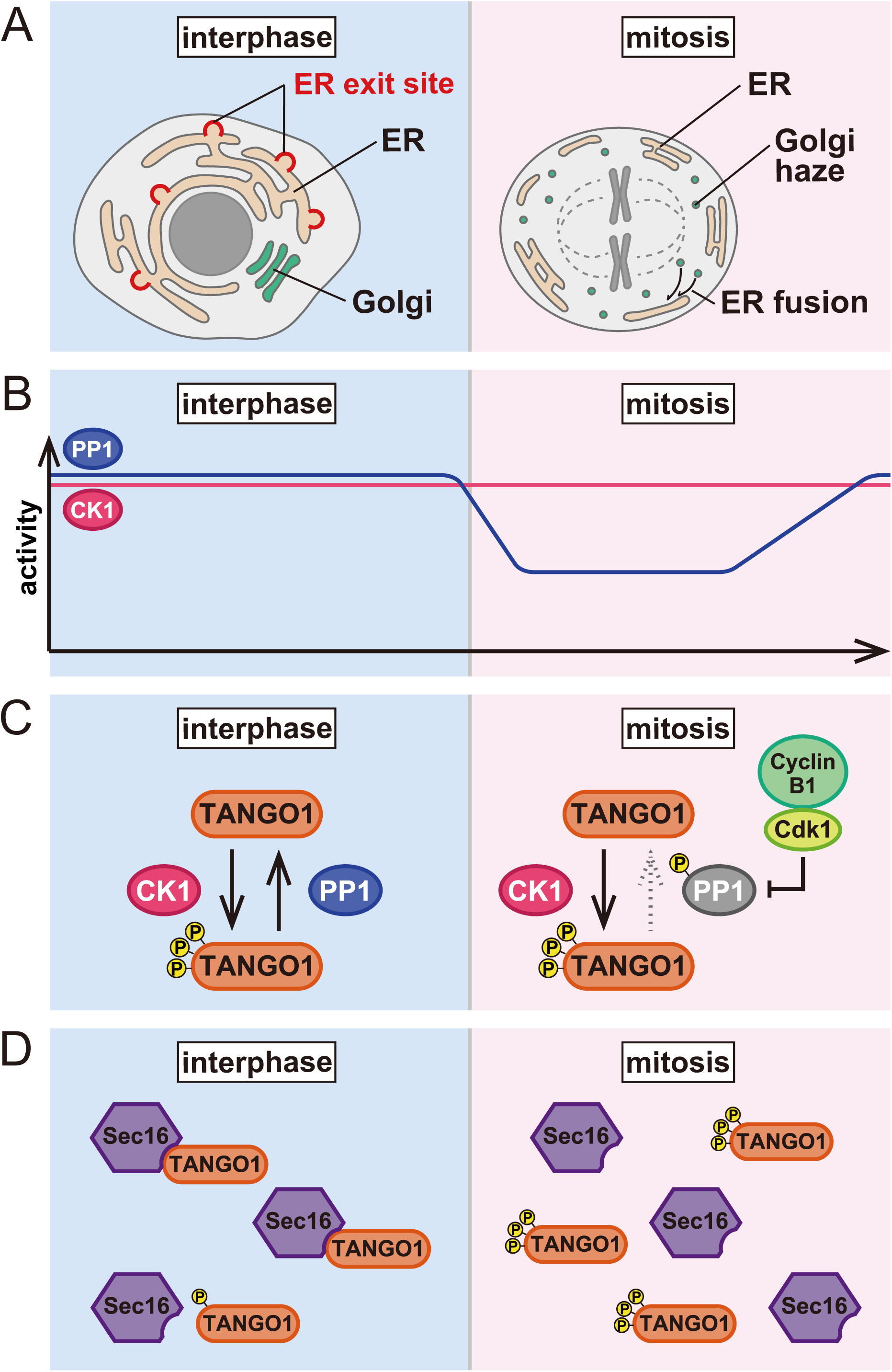
Schematic diagram of mitotic ER exit site dissociation. (A) ER and Golgi morphology during the cell cycle. In mitotic cells, Golgi membranes are fragmented and either stay as a Golgi haze or are fused with the ER. (B) Activity of CK1 and PP1 during the cell cycle. CK1 activity is unchanged throughout the cell cycle. Conversely, PP1 activity is suppressed during mitosis. (C) The phosphorylation status of TANGO1 is equilibrated via phosphorylation by CK1 and dephosphorylation by PP1 in interphase cells. During mitosis, the Cdk1/cyclinB1-dependent phosphorylation of PP1 reduces its activity and TANGO1 is hyperphosphorylated. (D) Interactions between TANGO1 and Sec16 result in ER exit site organization in interphase cells. The phosphorylation of TANGO1 reduces its binding to Sec16, leading to the dissociation of ER exit sites during mitosis.

We have discovered CK1 as a kinase that regulates TANGO1 PPS, and PP1 as a phosphatase for this domain. As shown in Fig. 8B, CK1 activity is rather constant throughout cell cycle progression, whereas PP1 activity is suppressed with phosphorylation by Cdk1–cyclin B1 at the onset of mitosis. The phosphorylation status of TANGO1 in interphase cells is somewhat at equilibrium based on phosphorylation by CK1 and dephosphorylation by PP1 (Fig. 8C). Conversely, TANGO1 is hyperphosphorylated during mitosis due to the loss of PP1 activity (Fig. 8C). We showed that TANGO1 with a phosphomimic mutant reduces interactions with Sec16, leading to the dissociation of ER exit sites (see Fig. 4A–D). Thus, we propose that ER exit site dissociation during mitosis is mediated by TANGO1 phosphorylation, which reduces binding to Sec16 (Fig. 8D). Indeed, our data suggest that overexpression of a non-phosphorylated form of TANGO1 inhibits ER exit site dissociation during mitosis (Fig. 5 F–H). Surprisingly, the ER exit site is restored by CK1 depletion in mitotic cells, suggesting that CK1 is a predominant kinase phosphorylating TANGO1. Our former study suggests that the depletion of TANGO1 (both TANGO1L and TANGO1S) results in the dissociation of ER exit sites (i.e. reduced colocalization rate between Sec16 and Sec31) and delays secretion. Thus, we speculate that mitotic TANGO1 phosphorylation also reduces secretion from the ER. Another COPII component, Sec24, is reportedly phosphorylated during mitosis, although the functional significance of this is not clear (Dudognon et al., 2004). We presume that a block in mitotic secretion would be achieved by the coordinated actions of COPII components regulated by mitotic kinases and phosphatases. TANGO1 should be in the center of these regulatory events.

Cells depleted of CK1 showed enlarged ER exit sites in interphase cells, strongly suggesting that phosphorylated TANGO1 exists to certain extent even in interphase cells (Fig. 8C). It is possible that TANGO1 phosphorylation is regulated not only by cell cycle progression but also by other cellular signaling. Indeed, it is known that CK1δ/ε regulate many cellular processes including the circadian rhythm (Xu et al., 2019; Yang et al., 2017). Thus, it would be interesting to determine whether TANGO1 phosphorylation by CK1 regulates ER exit site organization and secretion from the ER, which is dependent on cellular conditions such as circadian cycles. A yeast orthologue of CK1δ/ε, Hrr25, has been identified as a suppressor mutant for temperature-sensitive growth defects in the *Sec12-4* mutant (Murakami et al., 1999). Data indicate that Hrr25 negatively regulates ER export in yeast; however, the substrate responsible remains to be identified. TANGO1 is conserved only through metazoans; however, the function of CK1 in the negative regulation of ER export seems to be conserved from yeast to humans. CK1 also phosphorylates Sec23 to mediate COPII vesicle fusion with the Golgi (Lord et al., 2011). The phosphatase for Sec23 is PP6, and not PP1 (Bhandari et al., 2013). One interesting hypothesis would be that the phosphorylation status of these proteins might be regulated not by kinases but by phosphatases. In terms of regulatory activity, PP1 normally assembles to holoenzyme complexes with regulatory subunits. To date, hundreds of holoenzymes have been identified (Moura and Conde, 2019; Nilsson, 2019). Each holoenzyme shows distinct localization and different substrate binding properties according to the regulatory subunit. Thus, it is critical to identify the regulatory subunit responsible for TANGO1 dephosphorylation. Our current data did not preclude the possibility that TANGO1 PPS can be phosphorylated by kinases other than CK1 (and dephosphorylated by phosphatases other than PP1). It is worth determining whether the phosphorylation status of PPS can be regulated by external signals such as growth factors, cellular stress, and autophagy. Thus, TANGO1 might be a hub that controls ER exit site organization mediated by variations in cellular signaling.

In this study, we identified CK1δ/ε and PP1 as regulators of ER exit site organization during the cell cycle. In addition, we revealed that the phosphorylation status of TANGO1 is important for the regulation of ER exit site organization. This is the first report to describe the precise molecular mechanism of mitotic ER exit site dissociation.

## MATERIALS AND METHODS

### Antibodies

A female 6-wk-old Wistar rat (CLEA Japan) was immunized with FLAG-tagged Sec23A in TiterMax Gold (TiterMax USA). Splenocytes were fused with PAI mouse myeloma cells using polyethylene glycol (Roche). Hybridoma supernatants were screened by indirect ELISA with His-tagged Sec23A as the antigens. Positive hybridoma lines were subcloned, grown in serum-free medium (Nihon Pharmaceutical) supplemented with hypoxanthine-thymidine (Thermo Fisher Scientific), and purified with protein G-Sepharose (GE Healthcare; Saito et al., 2014). Polyclonal antibodies against Sec16-N (374–387 aa), Sec16-C (2,319–2,332 aa), and TANGO1-CT (1,884–1,898 aa for TANGO1L; 762–776 aa for TANGO1S) were raised in rabbits by immunization with keyhole limpet hemocyanin-conjugated peptides and affinity-purified by columns conjugated with the peptides (Thermo Fisher Scientific; Iinuma et al., 2007; Saito et al., 2009, 2011; Maeda et al., 2016; Tanabe et al., 2016). Polyclonal antibodies against Sec31 and TANGO1-CC1 were raised in rabbits by immunization with recombinant Sec31 (522-719 aa) or GST-tagged TANGO1L (1,231–1,340 aa) [TANGO1S (109–218 aa)] and affinity-purified by columns conjugated with GST-tagged Sec31 (522-719 aa) or ColdTF-tagged TANGO1 (1,231–1,340 aa) [TANGO1S (109–218 aa)] (Saito et al., 2011; Tanabe et al., 2016). Other antibodies were as follows: CK1δ (rabbit; Proteintech), CK1ε (mouse; BD), PPP1CA (mouse; Santa Cruz), PPP1CB (mouse; Santa Cruz), GAPDH (mouse; Merck), FLAG (mouse; Merck), FLAG (rat; agilent), HA (rat; Roche), and Sec31 (mouse; BD).

### Cell culture and transfection

HeLa and 293T cells were cultured in DMEM supplemented with 10% fetal bovine serum. Lipofectamine RNAi max (Thermo Fisher Scientific) was used for transfecting siRNA. For plasmids transfection, polyethylenimine “MAX” (polysciences) or Fugene 6 (Promega) were used. Doxycycline-inducible stable cell lines expressing TANGO1S-FLAG, TANGO1S (SA)-FLAG were made with lentivirus system described previously (Shin et al., 2006). Proteins were induced by incubation with 1 *μ*g/ml doxycycline for the indicated times.

### Double thymidine block

Cells treated with siRNAs were incubated for 22 h and then treated with 2.5 mM thymidine for 16 h. The cells were washed three times with PBS and released for 8 h. Then, the cells were incubated further with 2.5 mM thymidine for 16 h, washed three times with PBS and released for 10 h before fixation.

### Nocodazole block

Cells treated with siRNA were incubated for 59 h and then treated with 100 ng/ml nocodazole for 13 h before fixation or collecting by shake-off.

### Immunoprecipitation and western blotting

The experiments were essentially performed as described previously (Saito et al., 2014). Cells extracted with extraction buffer consisting of 20 mM Tris-HCl (pH7.4), 100 mM NaCl, 1 mM EDTA, 1% Triton X-100, and protease inhibitors were centrifuged at 20,000 g for 15 min at 4°C. Cell lysates were immunoprecipitated with FLAG M2 antibodies (SIGMA). The beads were washed with TBS/0.1% Triton X-100 for five times followed by elution with DYKDDDK peptide and processed for sample preparation.

### Biotinylated Phos-tag detection

Biotinylated Phos-tag analysis of phosphorylated proteins were conducted essentially as described previously (Kinoshita et al., 2006). The samples were resolved by SDS-PAGE followed by blotting to PVDF membranes. The membranes were probed with Phos-tag Biotin (Fuji Film) and then probed with Streptavidin-conjugated HRP (Fuji Film) for detection.

### siRNA oligos

Stealth select siRNAs for TANGO1S, TANGO1L, Sec23A, and Sec23B were purchased from Thermo Fisher Scientific. The oligo sequences used were

TANGO1S siRNA, 5’-GAAUUGUCGCUUGCGUUCAGCUGUU-3’; TANGO1L siRNA, 5’-CAACUCAGAGGAAAGUGAUAGUGUA-3’; Sec23A siRNA (1119), 5’-GGGUGAUUCUUUCAAUACUUCCUUA-3’; Sec23A siRNA (366), 5’-GCGUGGUCCUCAGAUGCCUUUGAUA-3’; Sec23B siRNA (1021), 5’-GCUGCAAAUGGUCACUGCAUUGAUA-3’; Sec23B siRNA (1923), 5’-CAGCAGCAUUCUAGCUGACAGAAUU-3’.

For control siRNA, Stealth RNAi™ siRNA negative control med GC (Thermo Fisher Scientific) was used.

Mission siRNA was purchased from Merck. The oligo sequences used were CK1δ siRNA (335), 5’-UGAUCAGUCGCAUCGAAUA-3’;

CK1δ siRNA (707), 5’-CCAUCGAAGUGUUGUGUAA -3’;

CK1ε siRNA (149), 5’-ACAUCGAGAGCAAGUUCUA-3’; CK1ε siRNA (734), 5’-CCUCCGAAUUCUCAACAUA-3’.

PPP1CA siRNA (423), 5’-GAGACGCUACAACAUCAAACUGUGG -3’.

PPP1CA siRNA (756), 5’-AGACGGCUACGAGUUCUUUGCCAAG -3’.

PPP1CB siRNA (563), 5’-UUAUGAGACCUACUGAUGU -3’.

PPP1CB siRNA (1137), 5’-CUAAUAGAAAGAUGUGCUACACUGU -3’.

For control siRNA, Mission siRNA Universal Negative Control #1 (Merck) was used. The number in the parentheses represents the starting base pair of the target sequence. Of note, PPP1CB siRNA (1137) is targeted to 3’UTR.

### Immunofluorescence microscopy

Immunofluorescence microscopy analysis was performed as described previously (Saito et al., 2014). Cells grown on coverslips were washed with PBS, fixed with methanol (6 min at −20°C), and then washed with PBS and blocked in blocking solution (5% BSA in PBS with 0.1% Triton X-100 for 30 min). After blocking, cells were stained with primary antibody (1 h at room temperature) followed by incubation with Alexa Fluor-conjugated secondary antibodies (Alexa Fluor 488, Alexa Fluor 568, and/or Alexa Fluor 647 for 1 h at room temperature). Images were acquired with confocal laser scanning microscopy (Plan Apochromat 63×/1.40 NA oil immersion objective lens; LSM 700; Carl Zeiss). The acquired images were processed with Zen 2009 software (Carl Zeiss). All imaging was performed at room temperature.

### Quantification of Pearson’s colocalization coefficient

HeLa cells were fixed and stained with indicated antibodies. Stained cells were analyzed by confocal laser scanning microscopy (Plan Apochromat 63×/1.40 NA oil immersion objective lens; LSM 700; Carl Zeiss) and processed with Zen 2011 software (Carl Zeiss). Intensity scanning and calculating coefficiency were performed by colocalization plugin Coloc2 in Fiji-ImageJ (Schindelin et al., 2012).

### Quantification of ER exit site size

HeLa cells were fixed and stained with indicated antibodies. Stained cells were analyzed by confocal laser scanning microscopy (Plan Apochromat 63×/1.40 NA oil immersion objective lens; LSM 700; Carl Zeiss) and processed with Zen 2011 software (Carl Zeiss). The size distribution of each ER exit site stained with Sec16 was analyzed by Fiji-ImageJ.

### In vitro kinase assay

0.57 *μ*M of purified recombinant GST-TANGO1 PPS+PRD-His_6_ (1,651–1,907 aa for TANGO1L; 529–785 aa for TANGO1S) or GST-TANGO1 PPS+PRD (SA)-His6 were incubated with 0.24 *μ*M of CK1δ (1–317aa), 20 mM Tris-HCl (pH 7.4), 150 mM NaCl, 0.1% Triton X-100, 11.4 mM MgCl2, and 4.75 mM [γ-P] ATP at 30 ℃ for 1h. Kinase assay was stopped by boiling with Laemmli Sample buffer. The samples were then subjected to SDS-PAGE followed by exposing with phosphor plate analyzed by Typhoon FLA9500 (GE healthcare).

### Online supplemental material

Figure S1 shows Sec23 depletion does not affect the enlargement of ER exit sites with IC261 treatment. Figure S2 shows purity of recombinant proteins used in the *in vitro* kinase assay. Figure S3 shows the degree of phosphorylation in TANGO1 PPS and its correlation with the dissociation of ER exit sites. Figure S4 shows that CK1 is required for mitotic ER exit site dissociation according to nocodazole block analysis.

## Acknowledgements

This work was supported by Grant-in-Aid for Scientific Research (18H06063 to M.M. and 17H03651, 19K22612 to K.S.) from the Ministry of Education, Culture, Sports, Science and Technology of Japan, by Takeda Science Foundation to K.S., The Uehara Memorial Foundation to K.S., and The Naito Foundation to K.S. and by Suzuken Memorial Foundation to M.M.

The authors declare no competing financial interests.

## Author contributions

Miharu Maeda designed and performed research, analyzed data, and wrote the manuscript; Yukie Komatsu performed research; and Kota Saito designed and performed research, analyzed data, and wrote the manuscript.

---

**Figure S1.**
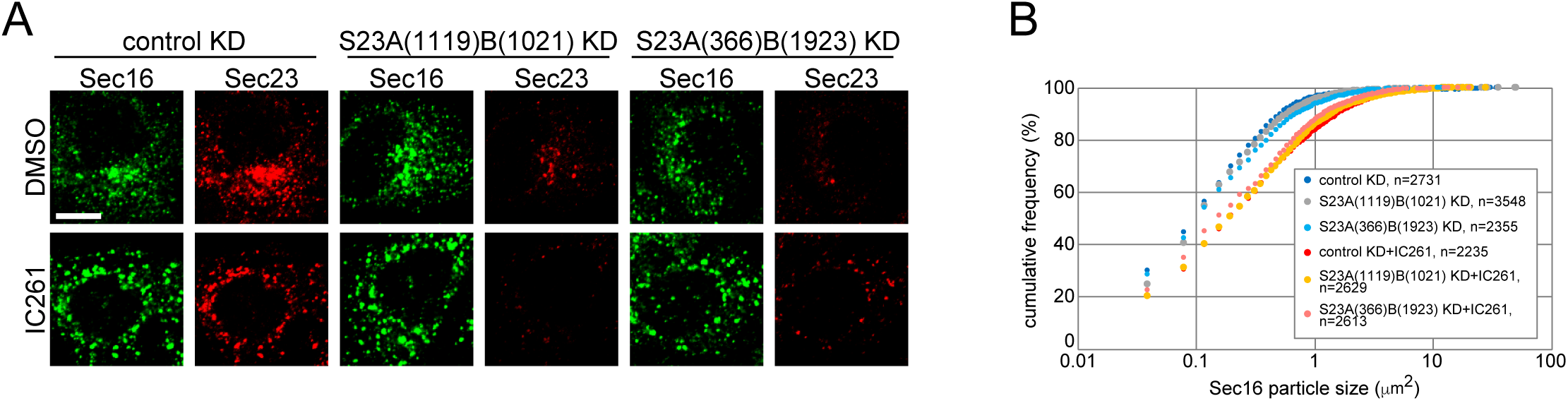
Sec23 depletion does not affect the enlargement of ER exit sites with IC261 treatment. (A) HeLa cells transfected with indicated siRNAs were treated with either DMSO or 10 *μ*M of IC261 for 12 h. Cells were fixed and stained with anti-Sec16-C and anti-Sec23 antibodies. Bars = 10 *μ*m. (B) Quantification of ER exit site size distribution from (A).

**Figure S2.**
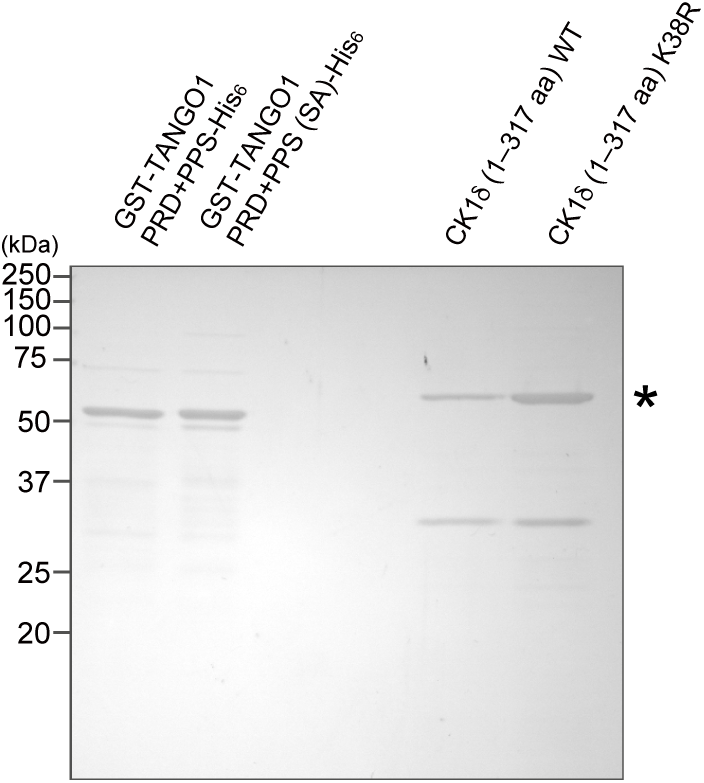
Recombinant proteins used in the *in vitro* kinase assay. Recombinant GST-TANGO1 PRD-His_6_, GST-TANGO1 PRD (SA)-His_6_, CK1δ (1–317 aa), and CK1δ KR (1–317 aa) were resolved by SDS-PAGE followed by Coomassie brilliant blue staining. The asterisks indicate bacterial protein contaminants.

**Figure S3.**
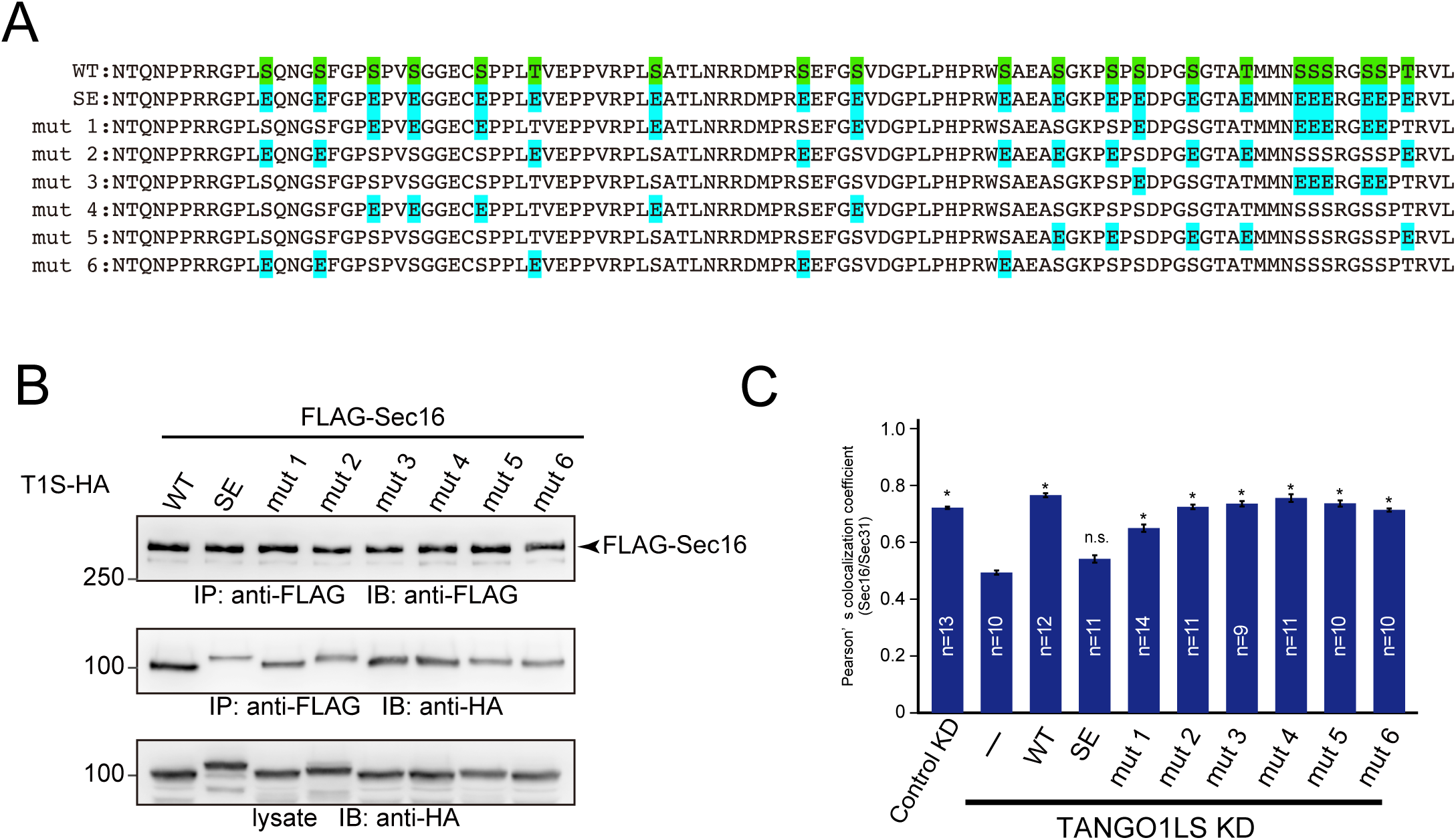
Degree of phosphorylation in TANGO1 PPS is correlated with ER exit site dissociation. (A) Sequences of human TANGO1 PPS mutants. (B) 293T cells were transfected with FLAG-Sec16 and the indicated TANGO1S-HA mutants. Cell lysates were immunoprecipitated with an anti-FLAG antibody and eluted with a FLAG peptide. Eluates were then subjected to SDS-PAGE followed by western blotting with anti-FLAG and anti-HA antibodies. (C) HeLa cells were treated with TANGO1L and TANGO1S siRNAs. After 24 h, the indicated TANGO1S-HA mutants were transfected and further cultured for 24 h. The cells were fixed and stained with anti-Sec16-C and anti-Sec31(m) antibodies and quantified based on Pearson’s colocalization coefficient. Error bars represent the means ± SEM. *p < 0.05 compared to TANGO1LS KD, n.s.: not significant compared to TANGO1LS KD.

**Figure S4.**
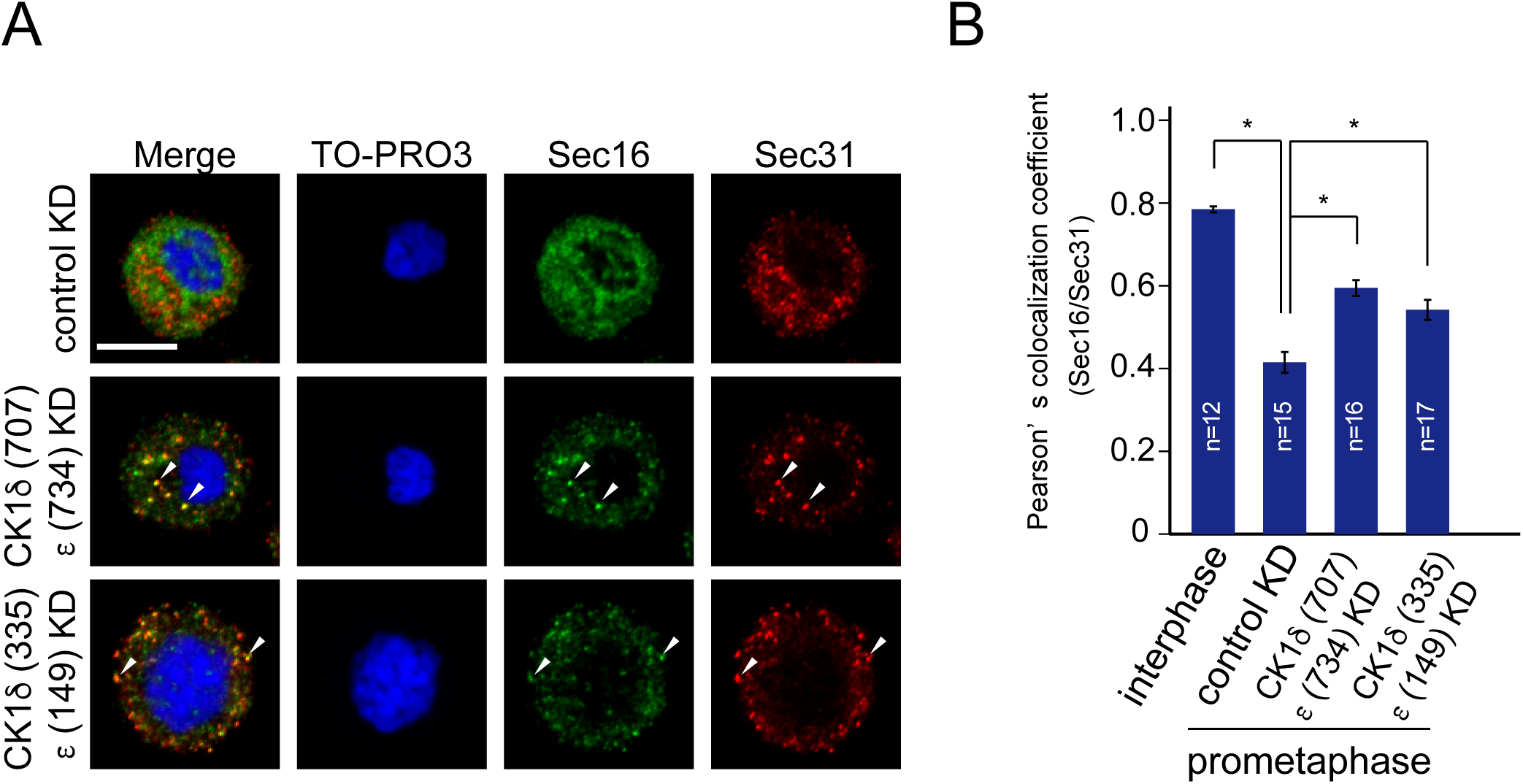
CK1 is required for ER exit site dissociation at prometaphase. (A) HeLa cells transfected with indicated siRNAs were synchronized in the mitotic phase by nocodazole block. Cells were then fixed and stained with anti-Sec16-C, anti-Sec31(m), and TO-PRO3. Bars = 10 *μ*m. (B) Quantification of Pearson’s colocalization coefficient for (A). Error bars represent the means ± SEM. *p < 0.05.

